# Butterfly and moth habitat specialisation changes along an elevational gradient of tropical forests on Mount Cameroon

**DOI:** 10.1101/2025.08.11.669622

**Authors:** Fernando P. Gaona, Sylvain Delabye, Vincent Maicher, Szabolcs Sáfián, Jan Altman, Jiří Doležal, Štěpán Janeček, Ishmeal N. Kobe, Mercy Murkwe, Robert Tropek

## Abstract

Niche breadth is a key ecological trait influencing species’ distribution, persistence, vulnerability to environmental change, and interspecific interactions. While the elevational niche-breadth hypothesis predicts broader ecological niches at higher elevations due to increased environmental stress and heterogeneity, empirical tests have mostly focused on food specialisation, with habitat specialisation remaining largely overlooked. Here, we provide the first direct test of this hypothesis using habitat characteristics, analysing fruit-feeding butterflies and moths along a complete tropical forest gradient on Mount Cameroon. We quantified habitat-related niche breadth using the Outlying Mean Index across 96 forest plots spanning 350–2200 m a.s.l., employing two complementary metrics: absolute niche breadth (ANB) based on the total range of habitat conditions occupied across the gradient, and within-elevation niche breadth (WENB) reflecting the proportion of local habitat heterogeneity utilized at each elevation. Forest structural heterogeneity increased steeply with elevation. In response, butterfly communities showed a substantial upslope broadening of ANB, particularly in Charaxinae and Satyrinae. Moth families exhibited similar but weaker patterns. In contrast, WENB showed no consistent elevational patterns and was more variable among lepidopteran groups. These results support the elevational niche-breadth hypothesis for habitat specialisation in tropical butterflies, likely reflecting increased habitat opportunity space and reduced competition at higher elevations. However, the decoupling of ANB and WENB highlights that broader availability of environmental conditions does not necessarily translate into more generalised local habitat use, suggesting constraints from behavioural preferences, resource distributions, or biotic interactions. By disentangling overall and local habitat niche breadths, our study reveals unrecognised complexity in elevational specialisation patterns and provides new insights into how tropical insect respond to environmental gradients.

## Introduction

Niche breadth, defined as the range of environmental conditions or resources a species can exploit, is a key ecological trait influencing how species interact with their environments and with other species in a community (MacArthur 1972; Devictor et al. 2010). Species with narrow niche breadths (specialists) are typically restricted to specific conditions or resources, often enabling efficient resource use but limiting their ability to tolerate heterogeneous or unstable environments. In contrast, generalists can persist across a broader range of habitats but may be less efficient or less competitive under stable conditions (Dennis et al. 2011).

Beyond the commonly studied specialisation for food resources, niche breadth can also reflect specialisation to habitat conditions, referring to the range of structural or microclimatic conditions a species occupies. Such habitat specialisation can be characterised as a partly flexible trait, responding to available abiotic conditions, their local heterogeneity, species interactions, and vegetation structure (Dennis et al. 2011; Rosenfeld 2003). Such habitat-related niche breadth plays a key role in the ecology and conservation of many groups, including herbivorous insects. In this group, spatial and structural patterns of habitat specialisation have been demonstrated, including vertical stratification of larvae and adult Lepidoptera (Basset et al. 2001; Delabye et al. 2021) or distinct herbivore communities in early versus late successional forests (Damken et al. 2012). Habitat specialisation also has strong ecological and conservation relevance, as habitat specialists tend to be generally more vulnerable to environmental change, particularly habitat loss and fragmentation (Nolte et al. 2019). For instance, most dung beetle species in the Amazon disappear from small forest fragments due to their limited habitat niches (Noble et al. 2023), and several beetle lineages in tropical mountains exhibit narrow elevational distributions tied to specific forest conditions (Macedo et al. 2017; Colares et al. 2021). Recent work has also shown that habitat specialisation is closely linked to dispersal ability, with habitat specialists exhibiting lower dispersal potential and thus higher sensitivity to resource instability and fragmentation (Habel et al. 2023).

Variation in niche breadth along environmental gradients has received substantial theoretical attention. Classical ecological theory predicts that stable and predictable environments favour specialisation, whereas broader niche breadths are advantageous under variable or stressful conditions, allowing species to persist despite changing resource availability or abiotic stress (MacArthur 1972; Dyer et al. 2007; Belmaker et al. 2012). While much of the theoretical and empirical attention has focused on latitudinal gradients, they are equally relevant to elevational gradients, where temperature, seasonality, and resource continuity often shift rapidly over short distances. These ideas have been formalised in the elevational niche-breadth hypothesis, which posits that species occurring at higher elevations should exhibit broader niche breadths due to environmental stress associated with lower temperatures, shorter growing seasons, and reduced resource availability (Rasmann et al. 2014).

Herbivorous insects, whose realised niches are shaped by abiotic conditions and their local heterogeneity, vegetation structure, and host plant distribution, are well suited for testing elevational patterns in niche breadth. However, virtually all existing studies have focused on food specialisation, typically quantified as dietary or host-plant niche breadth, while habitat-related specialisation has been almost entirely overlooked. Several studies have reported broader dietary niches at higher elevations in phytophagous beetles, bees, and butterflies in temperate mountains (Pradervand et al. 2013; Rasmann et al. 2014; Pellissier et al. 2012), with similar patterns observed in tropical bees in Tanzania (Classen et al. 2021) and moths in Ecuador and Costa Rica (Rodríguez-Castañeda et al. 2010). In contrast, no clear elevational trend was found in the host specificity of *Ficus*-associated moths in Papua New Guinea (Novotny et al. 2005). Whether herbivorous insects also shift their habitat specialisation along elevation, however, remains virtually unknown, particularly in tropical ecosystems.

This study addresses the lack of knowledge on elevational patterns of insect habitat specialisation by analysing habitat-related niche breadth of fruit-feeding butterflies and moths along the complete rainforest gradient of Mount Cameroon. We define habitat niche breadth as the range of compositional and structural forest conditions that individual species occupy. Fruit-feeding lepidopterans represent a suitable model group, as their adults can be sampled efficiently, and previous research has shown marked differences in habitat use between fruit-feeding butterflies and moths (e.g., Delabye et al. 2021). To capture different dimensions of habitat specialisation, we applied two complementary approaches: one assessing species’ habitat use relative to the variation in local conditions at each elevation, and the other measuring the total range of conditions occupied across the full gradient. These approaches allow us to distinguish behavioural specialisation from constraints imposed by environmental structure. Specifically, we test the hypothesis that lepidopteran communities exhibit broader habitat niches at higher elevations, as predicted by the elevational niche-breadth hypothesis (Rasmann et al. 2014). We also examine whether this pattern differs between butterflies and moths, as well as among their dominant taxonomic groups.

## Methods

### Lepidopteran datasets

We used a dataset of fruit-feeding butterflies and moths sampled along an Afrotropical elevational gradient on Mount Cameroon, originally published by Maicher et al. (2020a). It comprises 6,886 moth specimens representing 396 species and 16,508 butterfly specimens representing 121 species. Mount Cameroon (4.100° N, 9.050° E) is a tropical volcano rising directly from the Atlantic coast to 4,040 m a.s.l., hosting the only complete gradient of primary tropical rainforest from lowland to the natural timberline on the African continent (Maicher et al. 2020a). It is also a recognized biodiversity hotspot, particularly for butterflies and moths (e.g. Ustjuzhanin et al. 2018, 2025; Delabye et al. 2020). The region experiences a highly humid tropical climate, with mean annual rainfall exceeding 12,000 mm and a pronounced dry season from mid-November to February (Maicher et al. 2018, 2020a).

Butterflies and moths were sampled at six elevations (350, 650, 1100, 1500, 1850, and 2200 m a.s.l.), spanning lowland rainforest to montane forest. Each elevation included 16 plots (96 plots in total), each with a 25 m radius (Maicher et al. 2020a). Fruit-feeding Lepidoptera were captured using five baited traps per plot (four in the understory and one in the canopy), totalling 80 traps per elevation and 480 traps overall. Traps were baited with fermented mashed banana and operated for 10 consecutive days during each of three seasonal periods (wet–dry transition, dry– wet transition, and high dry season). All captured lepidopterans were removed daily, euthanised, and identified (see Maicher et al. 2020a for details). For analyses, specimens from all five traps and all three seasons were pooled per plot, and species were considered present in a plot only if more than four individuals were captured there.

### Niche breadth quantification

We quantified species’ habitat-related niche breadth (hereafter referred to as *niche breadth*) using the Outlying Mean Index (OMI; Dolédec et al. 2000), implemented via the *niche* function in the *ade4* package (Dray & Dufour, 2007). OMI is a multivariate ordination method that characterises species’ realised ecological niches in relation to environmental conditions across sampled sites. For each species, OMI calculates the deviation of its mean habitat from the overall habitat centroid and quantifies its niche breadth (termed *tolerance*) as the variance in habitat conditions across all its occurrences. Species with low tolerance values occupy narrower, more specialised niches, while those with high values exhibit broader, more generalised niches (Dolédec et al. 2000).

To analyse species’ habitat use, we used 27 habitat descriptors for each sampling plot related to forest structure and canopy openness, previously published by Delabye et al. (2021) (Table S1), which we standardised (mean = 0, SD = 1) prior to analyses. Based on these descriptors, we calculated two complementary indices of species’ niche breadth at each elevation, each capturing a distinct aspect of habitat specialisation. *Within-elevation niche breadth* (WENB) quantifies the proportion of the environmental variation available at a given elevation a species occupies, thus reflecting behavioural breadth relative to the local habitat mosaic. *Absolute niche breadth* (ANB), in contrast, is calculated in a single, gradient-wide environmental space and measures the raw physical span of habitat conditions a species occupies at a given elevation, regardless of local heterogeneity. Together, these metrics distinguish between the relative use of available habitat types (specialist vs. generalist behaviour) and the absolute environmental space that species inhabit.

Species’ within-elevation niche breadth (WENB) was quantified by running a separate principal component analysis (PCA) for each elevation using the habitat characteristics of the 16 plots sampled at that elevation and calculating tolerance for each species present. This approach allowed to express tolerance relative to the specific environmental heterogeneity present at each elevation. To make these tolerances comparable despite unequal habitat heterogeneity among elevations, we expressed each value as a percentage of the total inertia of the corresponding elevation-specific PCA, yielding the proportion of local environmental variation a species actually exploited. Species’ absolute niche breadth (ANB) was derived from a single PCA for all 96 plots, which established a common environmental frame across the entire gradient. Within this fixed frame, we calculated tolerance of each species separately at each elevation, based on its occurrence in the relevant subset of plots. These values represent the realised environmental envelope occupied by each species in standardised environmental units. Finally, we characterised each plot by community-weighted means (CWM) of both niche breadth indices, representing the average proportion of local habitat heterogeneity used by the local community (CWM WENB) and the average realised habitat breadth of the local community (CWM ANB).

### Analyses of elevational patterns

We analysed elevational patterns in niche breadth using generalised additive models (GAMs) fitted with the *mgcv* package (Wood 2017) in R 4.1.1 (R Core Team 2022). Although our hypothesis predicted linear responses, scatterplots suggested more complex relationships, for which penalised GAMs provide a single-step framework that retains full power for the linear case while remaining flexible enough to capture non-linear trends. For each response variable (CWM WENB and CWM ANB), we fitted a separate GAM with a penalised thin-plate regression spline of elevation (k = 10). The smoothing parameter was estimated using restricted maximum likelihood (REML), and a shrinkage penalty was included to allow the spline to collapse to a straight line when the effective degrees of freedom approach one, making the fit equivalent to a conventional GLM when supported by the data (Wood 2017). Model families were assigned after visual inspection of response histograms: Gaussian for approximately symmetric distributions, and Gamma otherwise (Table 1). The *gam*.*check* diagnostics confirmed that the selected basis dimension was adequate and that residuals showed no systematic pattern. To examine subgroup-specific patterns, we applied the same modelling framework to the three most abundant moth families (Erebidae, Noctuidae, and Geometridae) and the three most abundant butterfly subfamilies (Satyrinae, Limenitidinae, and Charaxinae). Plots lacking species of the focal (sub)family with the >4 individuals threshold were excluded from the corresponding taxon-specific analyses (Table 1).

**Table 1.**
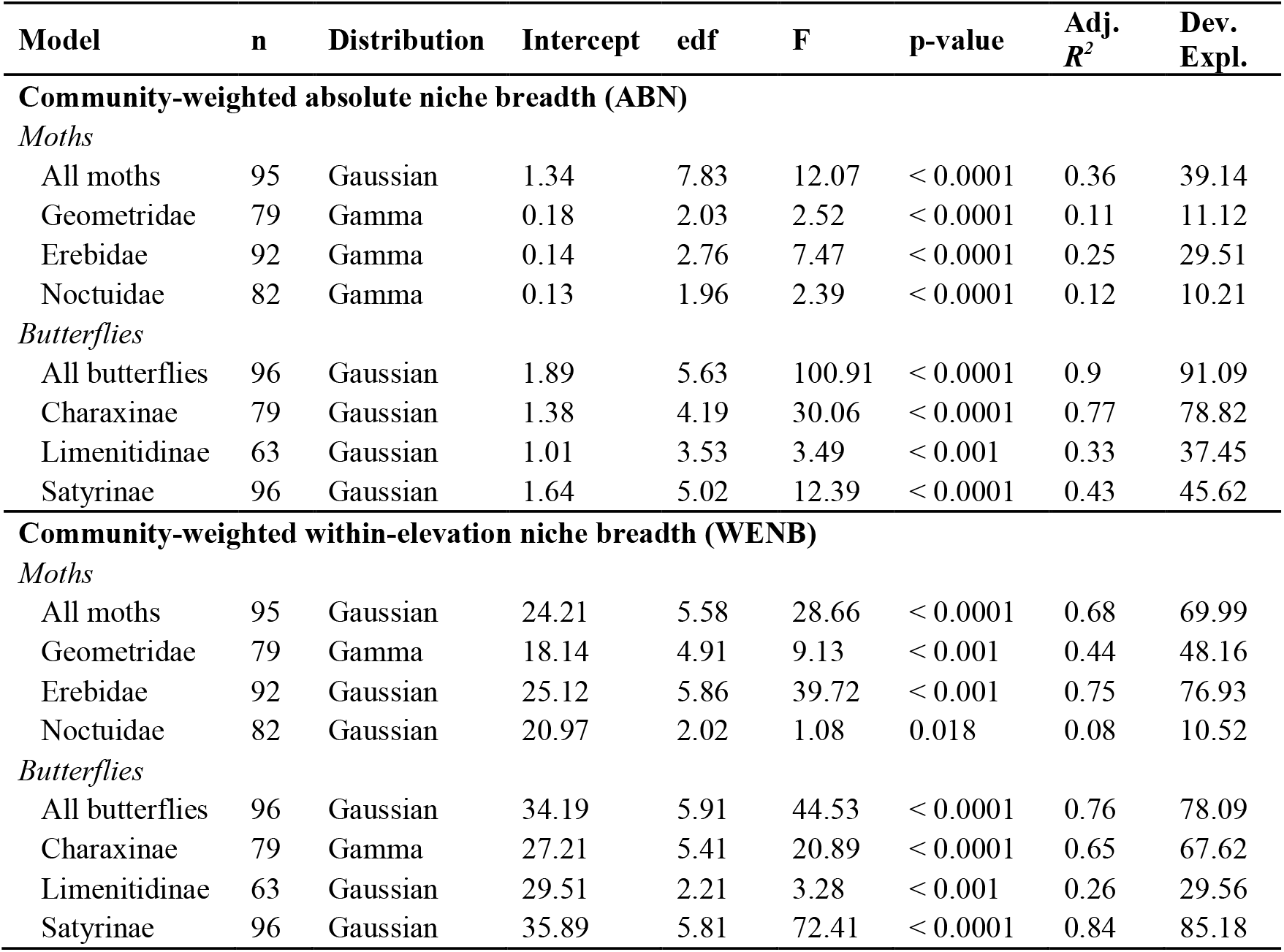
Results of generalised additive models (GAMs) testing the effect of elevation on community-weighted mean absolute niche breadth (ANB) and within-elevation niche breadth (WENB) of fruit-feeding Lepidoptera on Mount Cameroon. For each model (focal group), the table reports sample size (n, i.e. number of plots included), distribution family, model intercept, effective degrees of freedom of the smooth term (edf), and key model summary statistics. Sample sizes (n) vary because plots lacking any species of the focal group with ≥5-individuals were excluded from the respective analysis.

## Results

Habitat heterogeneity, represented by the total inertia of elevation-specific PCAs, increased strongly with elevation, rising more than seven-fold from the lowest sites to the upper mid-elevation (1850 m), before declining slightly at the highest elevation (2200 m; Table 2). This pattern indicates a pronounced upslope increase in structural variation of forest habitats along the elevational gradient, at least as captured by the 27 habitat characteristics used in this study.

**Table 2.**
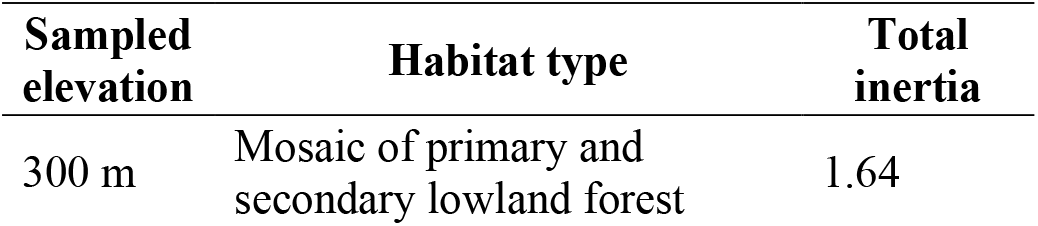
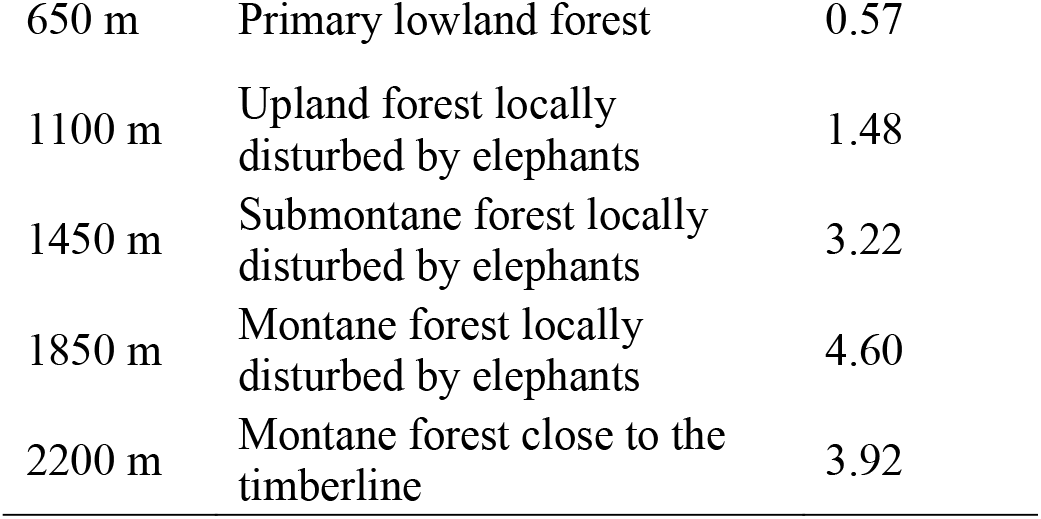
Habitat characterization and structural heterogeneity at each sampled elevation on Mount Cameroon. Heterogeneity is expressed as the total inertia from principal component analyses (PCA) of 27 habitat characteristics, with higher values indicating greater structural heterogeneity among the plots at a given elevation.

| Sampled elevation | Habitat type | Total inertia |
| --- | --- | --- |
| 300 m | Mosaic of primary and secondary lowland forest | 1.64 |
| 650 m | Primary lowland forest | 0.57 |
| 1100 m | Upland forest locally disturbed by elephants | 1.48 |
| 1450 m | Submontane forest locally disturbed by elephants | 3.22 |
| 1850 m | Montane forest locally disturbed by elephants | 4.60 |
| 2200 m | Montane forest close to the timberline | 3.92 |

All analyses of community-weighted mean absolute niche breadth (CWM ANB) showed significant relationships with elevation (Fig. 1; Table 1), though none followed a linear pattern. In moths, the full community showed no clear elevational pattern in CWM ANB (Fig. 1a), but each focal family exhibited a mild increase with elevation (Fig. 1b–d). In butterflies, the full community, as well as Charaxinae and Satyrinae, exhibited pronounced monotonic increases in CWM ANB with elevation, supported by relatively high adjusted *R*^*2*^ values (Fig. 1). Limenitidinae also showed a significant, though weaker, hump-shaped pattern (Fig. 1g). Overall, explained variation was higher in butterflies and their subgroups than in moths.

**Figure 1.**
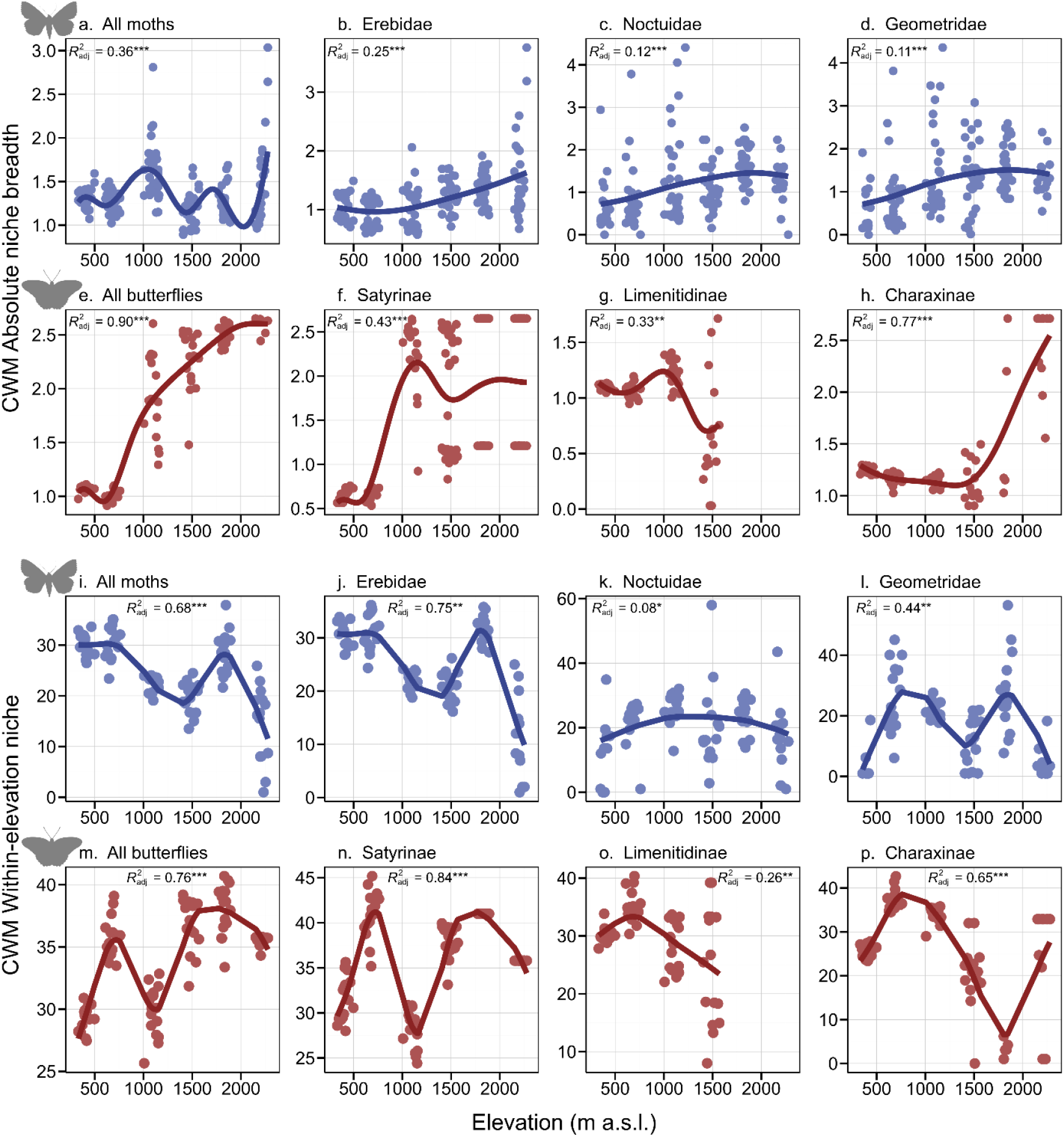
Elevational patterns in habitat-related niche breadth of fruit-feeding Lepidoptera on Mount Cameroon. Each panel shows the relationship between elevation and the community-weighted mean (CWM) of one of two niche breadth metrics: absolute niche breadth (ANB; panels a–h), calculated in a common environmental space across the entire gradient; and within-elevation niche breadth (WENB; i–p), expressed as a proportion of local habitat heterogeneity (see Methods for details). Blue symbols and lines represent moths (a–d, i–l), red symbols and lines represent butterflies (e–h, m–p). Trend lines represent Generalised Additive Models (GAMs), adjusted R^2^ values and the significance of the smooth term are shown in each panel (***p < 0.001, **p < 0.01, *p < 0.05).

All analyses of community-weighted within-elevation niche breadth (CWM WENB) also revealed significant relationships with elevation (Table 1), though none were linear. Compared to absolute niche breadth, elevational trends in CWM WENB were generally weaker, more complex, and more variable (Fig. 1i–p; Table 1). In moths, the full community and Erebidae exhibited non-monotonic patterns in CWM WENB, with generally decreasing trends interrupted by high values at upper mid-elevation plots (1850 m; Fig. 1i,j). Geometridae showed a bimodal pattern (Fig. 1l), while Noctuidae exhibited a modest mid-elevation peak (Fig. 1k). In butterflies, the full community displayed a gradual increase in CWM WENB with elevation, interrupted by high values at lower mid-elevation plots (650 m; Fig. 1m). Butterfly subfamilies showed contrasting responses, when Satyrinae displayed a bimodal pattern (Fig. 1n), Limenitidinae declined stadily with elevation (Fig. 1o), and Charaxinae showed a general decline in CWM WENB along elevation interrupted by high values at the highest elevation (2200 m; Fig. 1p). Explained variation in these models was generally lower than for CWM ANB, with strong fits observed only in a few studied groups (Table 1).

## Discussion

### Elevational niche-breadth patterns

Our study provides clear support for the elevational niche-breadth hypothesis (Rasmann et al. 2014) in the overall habitat niche breadth of fruit-feeding butterflies, but only limited support in fruit-feeding moths, indicating taxon-specific responses to increasing habitat heterogeneity (as also documented here) and harsher environmental conditions at higher elevations on Mount Cameroon. While full butterfly communities showed marked upslope broadening of habitat niches, with Charaxinae and Satyrinae displaying particularly consistent trends, moths exhibited no clear niche-breadth pattern in the full communities, though individual focal families showed mild increases. Interestingly, our complementary metric of local within-elevation habitat use revealed more complex and variable responses, with no consistent elevational trend in any studied group. These findings broadly align with previous studies reporting wider dietary niches at higher elevations in herbivorous insects, such as beetles, bees, and butterflies in both temperate (Pradervand et al. 2013; Pellissier et al. 2012; Rasmann et al. 2014) and tropical ecosystems (Rodríguez-Castañeda et al. 2010; Classen et al. 2021). However, to our knowledge, this is the first study testing the elevational niche-breadth hypothesis using habitat-related specialisation, rather than food specificity. Our results thus extend the generality of the hypothesis to habitat niches, at least for some groups of tropical insects.

Several general mechanisms may explain the observed upslope broadening of overall habitat niche breadth. The pronounced increase in forest structural heterogeneity with elevation likely provides a greater diversity of habitats and microhabitats, enabling species to occupy a broader range of conditions at higher elevations. Simultaneously, both lepidopteran abundance and species richness decline sharply with elevation on Mount Cameroon (Maicher et al. 2020a), which may reduce interspecific competition and facilitate the use of a broader range of available habitat patches. Harsher and more variable climatic conditions in montane forests may further favour broader ecological tolerances, consistent with the predictions of MacArthur (1972) and supported by similar findings in insect food specialisation (Pradervand et al. 2013; Pellissier et al. 2012; Rasmann et al. 2014; Rodríguez-Castañeda et al. 2010; Classen et al. 2021). Another contributing factor could be the scarcity or patchiness of key adult resources such as fruits and sap at higher elevations, potentially forcing individuals to exploit a wider range of habitat types, though this remains speculative due to the lack of data on resource distributions. Forest disturbance by elephants, which affects the gradient from mid-elevations upward (Table 2; Maicher et al. 2020a), may also enhance structural heterogeneity (Maicher et al. 2020b), potentially reinforcing upslope broadening of habitat niches, particularly in butterflies, which are more sensitive to tropical forest openness than moths (Delabye et al. 2021). Additionally, reduced predation pressure at higher elevations (e.g. Sam et al. 2014; Roslin et al. 2017) may allow butterflies to exploit more open or diverse microhabitats. Finally, repeated climatic oscillations over evolutionary timescales may have selected for broader environmental tolerances in montane insects (Janzen 1967; Ghalambor et al. 2006), although direct evidence for such evolutionary processes in Afrotropical insects is currently lacking. These mechanisms likely interact to shape the observed elevational patterns in overall habitat niche breadth, but their relative contributions remain uncertain and require further research.

### Overall niche breadth vs. local habitat use

The steep upslope increase in forest structural heterogeneity on Mount Cameroon raised diversity of habitat conditions at higher elevations, and fruit-feeding butterflies (and, to a lesser extent, some moth families) responded by broadening their overall habitat niche breadth (ANB). Yet this gradient produced no consistent pattern in within-elevation habitat use (WENB). This divergence indicates that tropical lepidopterans expand their niches by occupying a wider range of habitats when available along the gradient, but remain selective within elevations, using only a subset of the locally available conditions. The unclear and often erratic patterns in local habitat use may reflect limited behavioural flexibility or variable selectivity at local scales. Butterflies and moths are known to track specific microclimatic or microstructural conditions (Dennis et al. 2006; Shreeve 1992), but if these do not scale with overall structural heterogeneity, their local habitat use may not follow the upslope trend. Moreover, the erratic patterns may stem from the nature of the measured habitat characteristics, which describe forest structure and openness, whereas other key resources such as fermenting fruit, sap flows, or larval host plants may not follow the same elevational patterns. In such cases, lepidopteran communities may continue to use habitats highly selectively despite broader structural heterogeneity. Altogether, we suggest that the observed upslope broadening of overall habitat niche breadth likely reflects increased availability of diverse habitats, whereas local habitat use within elevations remains constrained by finer-scale ecological filters.

The elevational patterns in local habitat use showed a mid-elevation dip, relatively consistent among the studied lepidopteran groups, particularly around 1,100 m. This coincides with the mid-elevation peak in species richness, but not abundance, of both focal fruit-feeding lepidopteran groups on Mount Cameroon (Maicher et al. 2020a), suggesting intensified niche partitioning: when many species co-occur, competitive pressure may constrain each to a narrower subset of locally available habitat conditions (MacArthur 1972; Pianka 1974). Alternatively, key resources such as adult or larval food may peak at mid-elevations, allowing for finer specialisation and thus reduced their proportional habitat use, independent of overall habitat niche breadth. Although we cannot test this directly, tree diversity does not show a mid-elevation peak on Mount Cameroon (Hořák et al. 2019), implying that other resources, such as herbaceous plants or fruiting trees, may be more relevant. Overall, the mid-elevation dip in local habitat use likely reflects a combination of biotic interactions, unmeasured resource distributions, or other mechanisms, complicating interpretation of lepidopteran habitat specialisation along the elevational gradient.

## Conclusions

We provided the first direct test of the elevational niche-breadth hypothesis based on habitat characteristics in tropical herbivorous insects. Our results showed that overall habitat niche breadth increases with elevation in fruit-feeding butterflies and, to a lesser extent, in moths, supporting the idea that rising habitat heterogeneity at higher elevations expands the fundamental opportunity space available to species. However, this broader environmental range was not accompanied by broader within-elevation habitat use, likely due to constraints imposed by resource distribution, interspecific interactions, or behavioural preferences. By jointly analysing absolute and proportional habitat niche breadth, our study revealed a hidden complexity in elevational specialisation and highlighted the importance of disentangling these dimensions to better understand how insects respond to environmental gradients.

## Acknowledgements

The authors are grateful to everybody who helped with sampling and processing the previously published datasets. We used ChatGPT (models o3 and 4o; OpenAI) for English proofreading. Our research was authorized by multiple permits from the Ministries of the Republic of Cameroon for Forestry and Wildlife, and for Scientific Research and Innovation. This work was supported by the Czech Science Foundation.

## Data archiving statement

All data used in this study were already published in the Dryad repository, the data on lepidopteran communities are available in https://doi.org/10.5061/dryad.mgqnk98vr, and the data on habitat characteristics are available in https://doi.org/10.5061/dryad.vhhmgqnrm.

---

**Table S1.** Habitat descriptors and their brief definitions, recorded in 96 plots along an elevational gradient on Mount Cameroon and already used in Delabye et al. (2020).

| **Variable** | **Definition/way of evaluation** |
| --- | --- |
| *Forest structure* |  |
| Tree number | Number of all trees with >10 cm DBH (individuals/plot) |
| Live tree number | Number of live trees with >10 cm DBH (individuals/plot) |
| Dead tree number | Number of dead trees with >10 cm DBH (individuals/plot) |
| Tree basal area | Cumulative area of all tree trunks and stems with >10 cm DBH; calculated from measured DBH (m<sup>2</sup>) |
| Live tree basal area | Cumulative area of all live tree trunks and stems with >10 cm DBH; calculated from measured DBH (m<sup>2</sup>) |
| Dead tree basal area | Cumulative area of all dead tree trunks and stems with >10 cm DBH; calculated from measured DBH (m<sup>2</sup>) |
| Stand wood volume | Calculated from tree height and basal area of all trees (m<sup>3</sup>) |
| Live tree wood volume | Calculated from tree height and basal area of all living trees (m<sup>3</sup>) |
| Dead tree wood volume | Calculated from tree height and basal area of dead trees (m<sup>3</sup>) |
| Mean Diameter at Breast Height (DBH) | Calculated from DBH of all trees at the plot measured at 1.3 m above ground (cm) |
| Maximum Diameter at Breast Height | Maximum DBH from all trees at the plot (cm) |
| Mean tree height | Calculated from estimated heights of individual trees (m) |
| Maximum tree height | Estimated height of the highest tree at plot (m) |
| Mean stem slenderness Index | Ratio of estimated tree height to measured DBH foreach tree, averaged per plot |
| *Forest openness* |  |
| Maximum canopy openness | Maximum percentage of open sky from beneath a forest canopy determined from hemispherical photographs (%) |
| Mean canopy openness | Mean percentage of open sky from beneath a forest canopy determined from hemispherical photographs (%) |
| Herb layer cover | Herbs defined as all non-woody plants rooted in the soil; visually estimated over the entire plot area (%) |
| Shrub layer cover | Shrubs defined as all woody plants not exceeding the height of 4 m and with DBH <10 cm rooted in the soil and not classified as climbers; visually estimated over the entire plot area (%) |
| Herb and shrub layer cover | Combined cover of herb and shrub layers as described above (%) |
| Maximum Leaf Area Index | Maximum effective leaf area index integrated over the zenith angles 0° to 75° |
| Mean Leaf Area Index | Mean effective leaf area index integrated over the zenith angles 0° to 75° |
| Maximum amount of transmitted direct solar radiation transmitted by the canopy | Maximum amount of direct solar radiation transmitted by the canopy (%) |
| Mean transmitted direct solar radiation | Mean amount of direct solar radiation transmitted by the canopy (%) |
| Maximum amount of transmitted diffuse solar radiation | Maximum amount of diffuse solar radiation transmitted by the canopy (%) |
| Mean transmitted diffuse solar radiation | Mean amount of diffuse solar radiation transmitted by the canopy (%) |
| Maximum of total solar radiation | Maximum sum of direct and diffuse solar radiation transmitted by the canopy (%) |
| Mean total solar radiation | Mean sum of direct and diffuse solar radiation transmitted by the canopy (%) |

